# Danicamtiv reduces myosin’s working stroke but enhances contraction by activating the thin filament

**DOI:** 10.1101/2024.10.09.617269

**Authors:** Brent Scott, Lina Greenberg, Caterina Squarci, Kenneth S. Campbell, Michael J. Greenberg

**Author notes:** Corresponding author: Michael J. Greenberg Department of Biochemistry and Molecular Biophysics Washington University School of Medicine 660 S. Euclid Ave., Campus Box 8231 St. Louis, MO 63110.

## Abstract

Heart failure is a leading cause of death worldwide, and even with current treatments, the 5-year transplant-free survival rate is only ∼50-70%. As such, there is a need to develop new treatments for patients that improve survival and quality of life. Recently, there have been efforts to develop small molecules for heart failure that directly target components of the sarcomere, including cardiac myosin. One such molecule, danicamtiv, recently entered phase II clinical trials; however, its mechanism of action and direct effects on myosin’s mechanics and kinetics are not well understood. Using optical trapping techniques, stopped flow transient kinetics, and *in vitro* reconstitution assays, we found that danicamtiv reduces the size of cardiac myosin’s working stroke, and in contrast to studies in muscle fibers, we found that it does not affect actomyosin detachment kinetics at the level of individual crossbridges. We demonstrate that danicamtiv accelerates actomyosin association kinetics, leading to increased recruitment of myosin crossbridges and subsequent thin filament activation at physiologically-relevant calcium concentrations. Finally, we computationally model how the observed changes in mechanics and kinetics at the level of single crossbridges contribute to increased cardiac contraction and improved diastolic function compared to the related myotrope, omecamtiv mecarbil. Taken together, our results have important implications for the design of new sarcomeric-targeting compounds for heart failure.

**SIGNIFICANCE STATEMENT:** Heart failure is a leading cause of death worldwide, and there is a need to develop new treatments that improve outcomes for patients. Recently, the myosin-binding small molecule danicamtiv entered clinical trials for heart failure; however, its mechanism at the level of single myosin crossbridges is not well understood. We determined the molecular mechanism of danicamtiv and showed how drug-induced molecular changes can mechanistically increase heart contraction. Moreover, we demonstrate fundamental differences between danicamtiv and the related myosin-binding small molecule omecamtiv mecarbil that explain the improved diastolic function seen with danicamtiv. Our results have important implications for the design of new therapeutics for heart failure.

## INTRODUCTION

Heart failure, a leading cause of mortality and morbidity in the world, is characterized by the inability of the heart to generate sufficient power to perfuse the body at normal filling pressures. Heart failure with reduced ejection fraction (HFrEF) is characterized by reduced contractility during systole, and it accounts for ∼50% of heart failure cases (1). Current treatments for HFrEF (e.g., beta blockers and ACE inhibitors) target adverse remodeling of the heart, which occurs secondary to reduced contractile function. While targeting adverse remodeling has significantly improved outcomes for patients with HFrEF, the 5-year transplant-free survival rates are still only ∼50-70% (2). Thus, there is an outstanding need to develop new therapeutics that improve mortality and quality of life for patients with HFrEF. There has been a long-standing interest in developing heart failure treatments that reverse the reduced contractile function seen in patients with HFrEF, but this approach has been met with several challenges. Inotropes, such as milronone, increase heart contractility by modulating calcium flux; however, elevated calcium leads to increased mortality (3), and as such, inotropes are typically only used with patients in end-stage heart failure (1). Recently, there have been several efforts to develop small molecules that directly target the sarcomeric machinery to increase cardiac contraction without affecting calcium handling (4–9). The first of these drugs was omecamtiv mecarbil (OM) (4), which was discovered in a high-throughput screen for small molecules that increase cardiac myosin’s ATPase activity. OM made it to phase III clinical trials (10); however, the FDA declined to approve OM due to its limited effect size and negative effects on relaxation.

Recently, a new myosin-binding compound was reported, danicamtiv, and this small molecule is currently in phase II clinical trials for HFrEF (11). Currently, there is limited information about the mechanism of danicamtiv, and the direct molecular effects of danicamtiv on the kinetics and mechanics of individual myosin crossbridges is poorly understood. In a study comparing OM and danicamtiv, it was reported that danicamtiv has a smaller impact on relaxation compared to OM (9); however, the molecular mechanism underlying this difference is not known. Moreover, elegant experiments in muscle fibers and myofibrils have shown clear structural and functional effects of danicamtiv at the cellular and sub-cellular levels (12–14). X-ray diffraction of danicamtiv treated muscle fibers revealed an increase in filament lattice spacing and a re-positioning of myosin heads closer towards the thin filament (14). Moreover, danicamtiv was shown to slow the rate of tension development in porcine myofibrils (14), slow tension redevelopment in human myocardial bundles (13), slow the rate of myofibril relaxation (14), and slow the rate of myosin-driven motility *in vitro* (14). Based on these observations, it was suggested that danicamtiv likely slows the rate of ADP release from actomyosin, the transition that limits actomyosin dissociation in cycling muscle; however, the direct effects of danicamtiv at the level of single crossbridges has not been examined.

Here, we use a combination of stopped flow transient kinetics, single molecule optical trapping, computational modeling, and in vitro reconstitution assays to directly measure how danicamtiv affects the mechanics and kinetics of individual myosin crossbridges. Our results provide new insights into the mechanism of danicamtiv and help to explain its differences with OM.

## RESULTS

We set out to measure the effects of danicamtiv on the mechanics and kinetics of individual cardiac myosin crossbridges. We used porcine cardiac actin, which is identical to the human isoform, and porcine ventricular cardiac myosin which has biophysical properties that are indistinguishable from human cardiac myosin (15–17). Danicamtiv was dissolved in DMSO (10 mM) and diluted in KMg25 Buffer to a final concentration of 10 µM for all experiments. The final experiment buffers contained 0.1% DMSO. First, we examined the effects of danicamtiv on myosin’s steady-state ATPase activity. Similar to previous reports using isolated myofibrils (11), we found that danicamtiv increases myosin’s maximal steady-state ATPase rate at saturating actin by ∼1.2-fold (P = 0.028) and increases the effective K_m_ (P = 0.01) (**Fig. 1A**, **Table 1**). Next, we used an *in vitro* motility assay in which fluorescently labeled actin is translocated over a bed of myosin in the presence of ATP (18) and we measured the speed of myosin-based movement. Consistent with previous reports, we observed that danicamtiv reduces the motile speed by ∼55% (**Fig. 1B**) (P = 4×10^-6^). Taken together, danicamtiv accelerates the overall rate of myosin crossbridge cycling kinetics while reducing the speed at which it moves actin.

**Figure 1.**
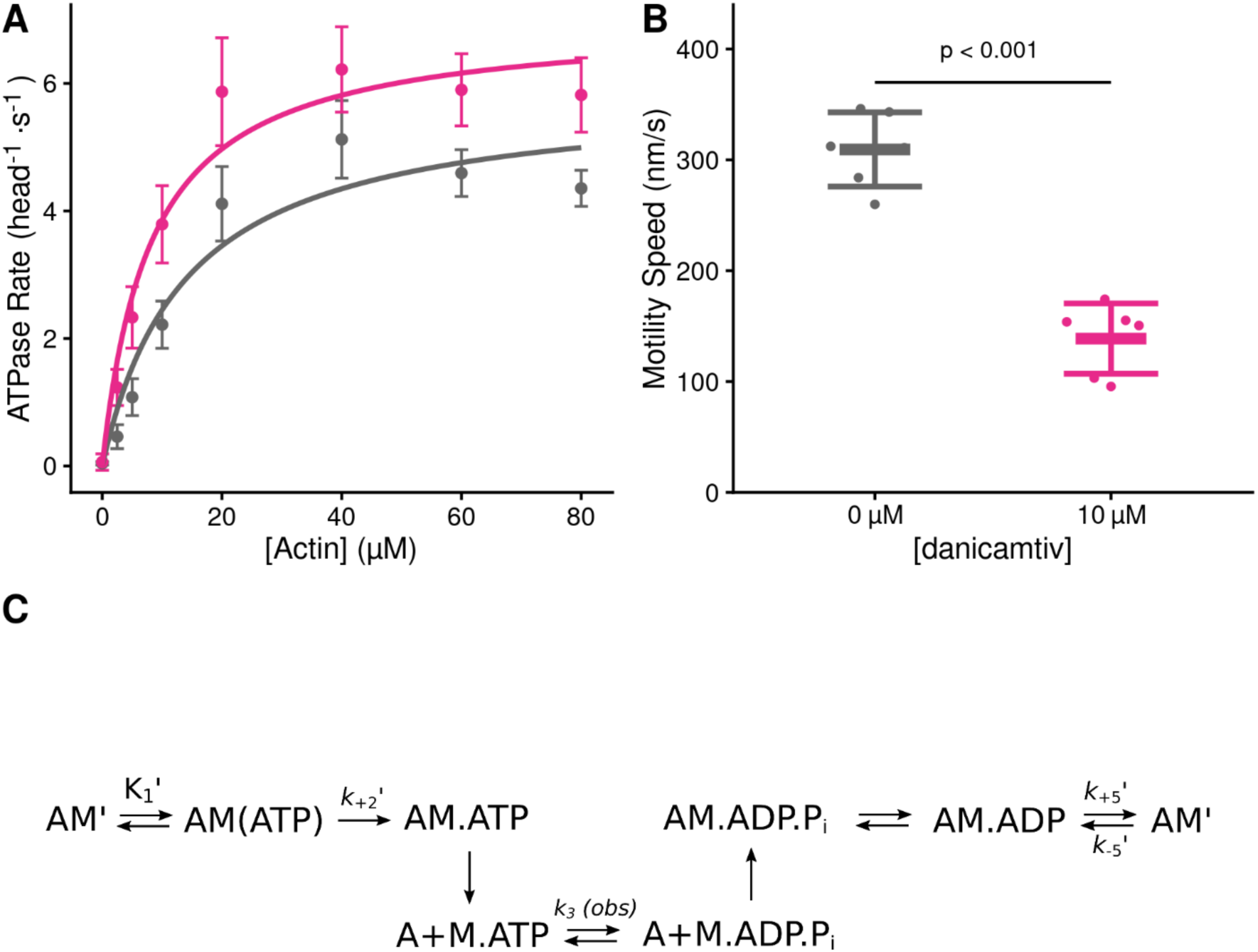
Steady state properties of β-cardiac myosin treated with danicamtiv. **A)** The steady-state myosin ATPase rate was measured using the NADH coupled assay. The steady state myosin ATPase rate is plotted verses a function of actin concentration. Data were fitted by a hyperbolic function to calculate the maximal cycling rate (V_max_) and actin affinity (K_m_) with the Michaelis Menten equation. Treatment with 10 µM danicamtiv increased the maximal rate by ∼1.2 fold from 5.9 to 7.0 s^-1^ (P = 0.028) and decreased the K_m_ from 14.0 to 8.1 µM (P = 0.01). Each point represents the average rate from four independent trials with error bars showing the standard deviation. Statistical testing done using a 2-tailed T-test. Black = DMSO control. Pink = 10 µM danicamtiv. **B)** Speed of actin translocation in the unregulated in vitro motility assay. The addition of 10 µM danicamtiv decreased motility speed ∼55% (P = 4×10^-6^). Thick horizontal lines show the average speed with standard deviation shown by the thin horizontal lines. Points represent the average speed of all filaments in a field of view for a single technical replicate measured across N = 3 independent experiments. Statistical testing done using a 2-tailed T-test. **C)** Scheme of myosin’s mechanochemical cross-bridge cycle. Myosin’s rate limiting step is actin attachment, so the predominant population of motors reside in the pre-working stroke M.ADP.Pi state during steady state cycling. The steady-state ATPase is thus limited by actin attachment (*k_att_*) which is rapidly followed by the mechanical working stroke and phosphate release. *In vitro* motility speed is limited by the ADP release rate (*k_+5_*’).

**Table 1.** Solution kinetics summary.

| Parameter | 0 $\mu\text{M}$<br>danicamtiv | 10 $\mu\text{M}$<br>danicamtiv | P-value |
| --- | --- | --- | --- |
| **Steady State ATPase** |  |  |  |
| $V_{\text{max}}$ ($\text{head}^{-1} \cdot \text{s}^{-1}$) | $5.9 \pm 0.4$ | $7.0 \pm 0.6$ | 0.028 |
| $K_{\text{m}}$ ($\mu\text{M}$) | $14.0 \pm 2.5$ | $8.1 \pm 1.6$ | 0.01 |
| Catalytic Efficiency [$V_{\text{max}}/K_{\text{m}}$] ($\mu\text{M}^{-1} \cdot \text{s}^{-1}$)* | $0.42 \pm 0.08$ | $0.86 \pm 0.19$ | 0.013 |
| **ATP Induced Dissociation** |  |  |  |
| $1/K_1'$ ($\mu\text{M}$) | $269 \pm 71$ | $245 \pm 38$ | 0.64 |
| $k_{+2}'$ ($\text{s}^{-1}$) | $1054 \pm 123$ | $1053 \pm 86$ | 0.99 |
| $K_1' \cdot k_{+2}'$ ($\mu\text{M}^{-1} \cdot \text{s}^{-1}$)* | $4.0 \pm 0.7$ | $4.4 \pm 1.1$ | 0.64 |
| **ADP Release** |  |  |  |
| $k_{+5}'$ ($\text{s}^{-1}$) | $73.2 \pm 3.3$ | $73.1 \pm 3.3$ | 0.96 |
| $k_{-5}'$ ($\mu\text{M}^{-1} \cdot \text{s}^{-1}$)* | $4.1 \pm 0.3$ | $3.9 \pm 0.2$ | 0.40 |
| $K_5'$ ($\mu\text{M}$) | $18.0 \pm 4.6$ | $18.6 \pm 4.1$ | 0.87 |
| **ATP Hydrolysis** |  |  |  |
| $k_3$ (*obs*) ($\text{s}^{-1}$) | $79.8 \pm 3.0$ | $72.6 \pm 2.5$ | 0.03 |
| **Single turnover assay** |  |  |  |
| $k_{\text{det}}$ ($\mu\text{M}^{-1} \cdot \text{s}^{-1}$) | $3.8 \pm 0.9$ | $4.2 \pm 0.8$ | 0.23 |
| $k_{\text{att}}$ ($\mu\text{M}^{-1} \cdot \text{s}^{-1}$) | $0.0040 \pm 0.0002$ | $0.0063 \pm 0.0007$ | < 0.001 |
| *calculated value |  |  |  |

### Danicamtiv does not affect key biochemical transitions that govern actomyosin detachment (ADP release or ATP-induced dissociation)

Our data demonstrate that danicamtiv increases overall cycling kinetics while simultaneously reducing the speed of myosin motility, suggesting that danicamtiv affects the coupling between the mechanics and kinetics of myosin crossbridges. In the motility assay, the speed of actin filament translocation is proportional to the displacement generated by a single myosin (i.e., the size of the myosin working stroke) divided by the amount of time that myosin remains attached to the actin filament (19):

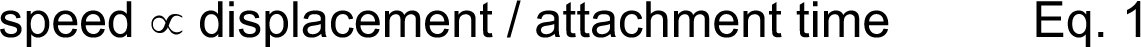

Thus, the ∼50% reduction in motile speed suggests that either the displacement generated by the myosin is reduced by half or the attachment time of a crossbridge gets twice as long. Therefore, we set out to directly test these two possibilities.

To test whether danicamtiv affects the time that crossbridges remain bound to actin, we measured the rates of key biochemical transitions. The amount of time that actin and myosin remain attached is set by the time required for ADP release and subsequent ATP-induced actomyosin dissociation (20). Since the rate of a transition is inversely proportional to the average time for the transition to occur, we can equivalently state that the rate of crossbridge dissociation is set by the rates of ADP release from myosin and ATP-induced dissociation of actomyosin (**Fig. 1C**). Therefore, we measured the rates of these transitions using stopped flow transient kinetics.

First, we measured the rate of ATP-induced actomyosin dissociation using pyrene-labeled actin as a fluorescent reporter of actomyosin binding (21), where myosin detachment from pyrene-labeled actin causes an increase in pyrene fluorescence. We rapidly mixed pyrene-labeled actomyosin with a range of ATP concentrations and measured the change in fluorescence. As previously described, fluorescence transients were well fitted by the sum of two exponential functions, where the observed rate of the fast phase can be used to report the rate of ATP-induced actomyosin dissociation (21). The relationship between the observed rate of the fast phase and the concentration of ATP was fitted with a hyperbolic function to obtain the maximal rate of ATP-induced dissociation at saturating ATP concentrations, k_+2_’, and the concentration of ATP necessary to reach half-maximal saturation, 1/K_1_’ (**Fig. 2A**). We found that danicamtiv does not affect the maximal rate of ATP-induced dissociation (P = 0.99), the concentration of ATP necessary to reach half-maximal saturation (P = 0.64), or the second-order rate of ATP induced dissociation (4.0 ± 0.7 vs 4.4 ± 1.1 µM^-1^s^-1^, for 0 and 10 µM danicamtiv, P = 0.64) (**Table 1**). Thus, changes in ATP-induced actomyosin dissociation cannot explain the reduced motility seen with danicamtiv.

**Figure 2.**
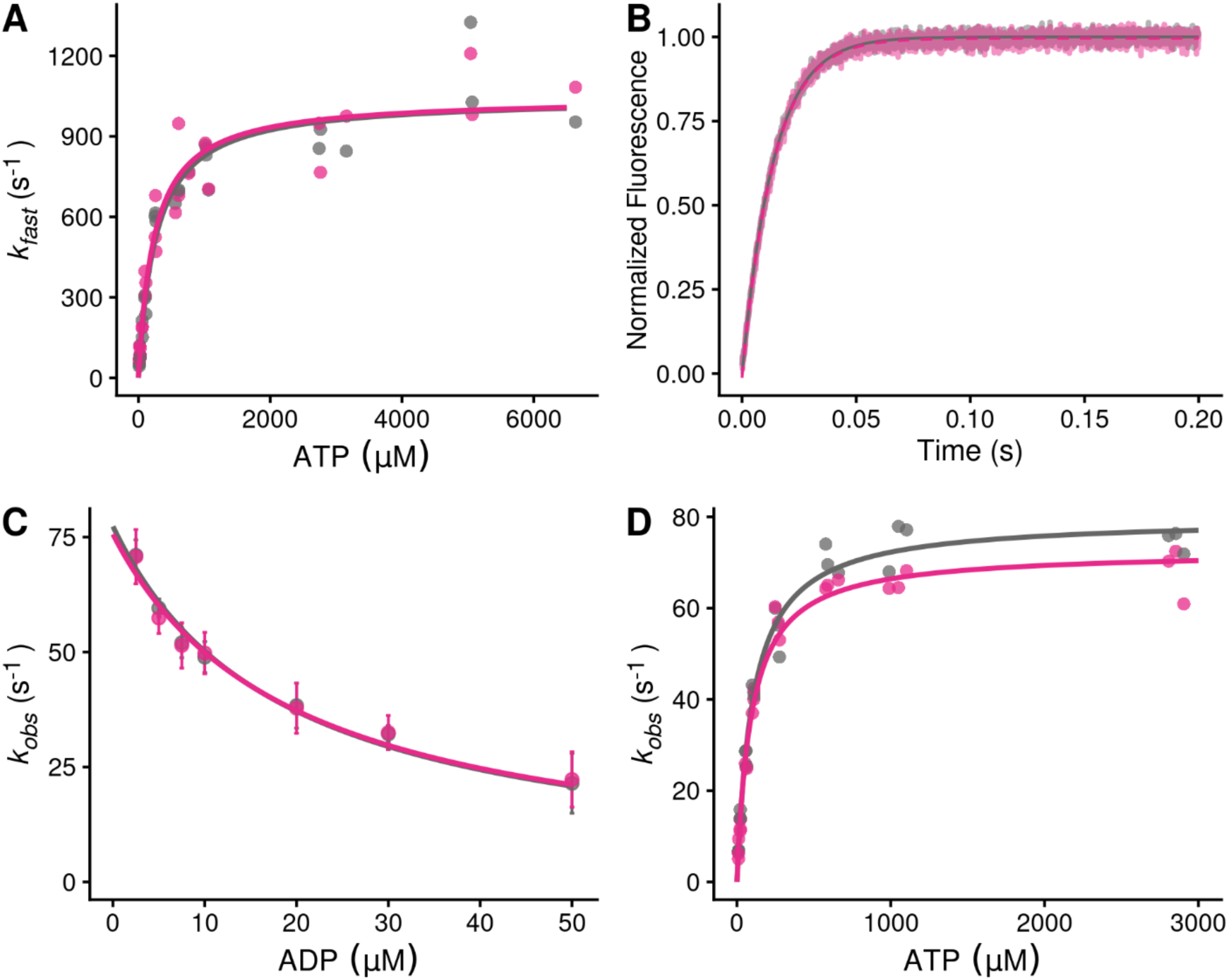
Stopped-flow kinetics measured with and without 10 µM danicamtiv. **Black** = DMSO control. Pink = 10 µM danicamtiv. **A)** The rates of ATP-induced actomyosin dissociation were measured in the stopped flow. Transients were well fitted by the sum of two exponential functions, where the observed rate of the fast phase (*k_fast_*) is plotted as a function of ATP concentration. Data were fitted with a hyperbolic function to obtain K_1_’ and k_+2_’. There are no differences in either K_1_’ or k_+2_’ with or without danicamtiv (P = 0.64 and P = 0.99, respectively). Each point represents an independent measurement over 3 experimental days. **B)** The rate of ADP release from actomyosin was measured using stopped flow techniques, and the fluorescence transients were fitted with single exponential functions. Note, both 0 and 10 µM danicamtiv are plotted and overlay. There is no statistically significant difference in the rate of ADP release the absence or presence of danicamtiv (P = 0.96). **C)** The overall ADP binding affinity to actomyosin was measured by mixing an actomyosin solution containing increasing concentrations of ADP with 50 µM ATP (concentrations after mixing) measured using a competition experiment. The observed rate as a function of ADP concentration was fitted with a hyperbolic function to determine the ADP affinity (see methods). Each point shows the average of 3 separate trials and error bars show the standard deviations. There is no difference in the ADP binding affinity (*k_-5_’*; P = 0.40). **D)** The rate of ATP hydrolysis by myosin was measured using stopped flow techniques. The rate of hydrolysis is reported by the change in tryptophan fluorescence at saturating ATP concentrations. Fluorescence transients are well fitted with single exponential functions. The observed rates of ATP hydrolysis were plotted against their respective ATP concentration and fitted with a hyperbolic function. The plateau represents the sum of the forwards and backwards rate of ATP hydrolysis (*k_3_ (obs)*). While there was a slight decrease in the observed hydrolysis rate with danicamtiv (P = 0.03), this slight decrease is not biologically meaningful. For all stopped-flow values, see **Table 1**.

Next, we measured the rate of ADP release from actomyosin by preparing a mixture of ADP-saturated myosin and pyrene-labeled actin, and rapidly mixing this with saturating amounts of ATP, leading to an increase in fluorescence (21). The fluorescence transients were fitted with single exponential functions to obtain the rate of ADP release from actomyosin (**Fig. 2B**). We found that the rates of ADP release with and without 10 µM danicamtiv were not statistically different (WT: 73 ± 3 s^-1^, Danicamtiv: 73 ± 3 s^-1^, P = 0.96). This rate of ADP release is consistent with previous measurements using porcine β-cardiac myosin (16, 22) and recombinant human cardiac myosin (23). Moreover, at physiologically relevant saturating ATP concentrations, the rate of ADP release is slower than the rate of ATP-induced actomyosin dissociation both in the presence and absence of danicamtiv. As such, the rate of ADP release limits actomyosin dissociation at saturating ATP concentrations, such as those in the motility assay and in cardiomyocytes. Taken together, the observed reduction in motile speed cannot be explained by changes in the rate of ADP release from actomyosin.

It has also been proposed that danicamtiv might affect the rate of ADP binding to actomyosin (14). To test this, we determined the ADP binding affinity to actomyosin by measuring the rate of pyrene-actomyosin dissociation in the presence of competing mixtures of ATP and ADP (21). We found that the ADP affinity was not statistically different with or without danicamtiv (**Fig. 2C**; P = 0.87). Moreover, we can use this affinity and the measured ADP release rate to calculate the rate of ADP binding (*k_-5_’* see Methods for details), and we found that there is no statistical difference in the rate of ADP rebinding with danicamtiv (P = 0.4). Taken together, we did not observe changes in the rates of key steps of actomyosin dissociation that could explain the reduction in motile speed.

Finally, we measured the rate of ATP hydrolysis by myosin across a range of ATP concentrations using the intrinsic tryptophan fluorescence of the myosin which increases with hydrolysis (**Fig. 2D**) (21). We found that the observed fluorescence transients are well fitted by a single exponential function. When plotting the observed rates against their respective ATP concentrations, the data is well described by a hyperbolic function where the plateau represents the sum of the forwards and backwards rates of ATP hydrolysis. Our measured rate of ATP hydrolysis by porcine cardiac myosin (79.8 ± 3.0 s^-1^) is consistent with previous reports (23). While there was a slight decrease in the observed hydrolysis rate with danicamtiv (79.8 ± 3.0 vs. 72.6 ± 2.5 s^-1^, for DMSO control and danicamtiv, respectively; P = 0.03), this small decrease is not biologically meaningful, and it cannot not explain the differences see with the ATPase rate or motility.

### Danicamtiv reduces the displacement of myosin’s working stroke without altering detachment kinetics

Given that we did not observe a change in biochemical kinetics that could explain the ∼50% reduction in motile speed with danicamtiv (**Fig. 1B**), we used single molecule optical trapping techniques to measure the mechanics of the cardiac myosin working stroke in the presence and absence of danicamtiv (16, 24). We used the three-bead assay in which an actin filament is suspended between two optically-trapped beads and lowered onto a surface-bound pedestal that is sparsely coated with myosin (25) (**Fig. 3A**). We were able to clearly resolve single molecule interactions between actin and myosin (**Fig. 3B**), enabling us to probe the mechanics and kinetics of the myosin working stroke.

**Figure 3.**
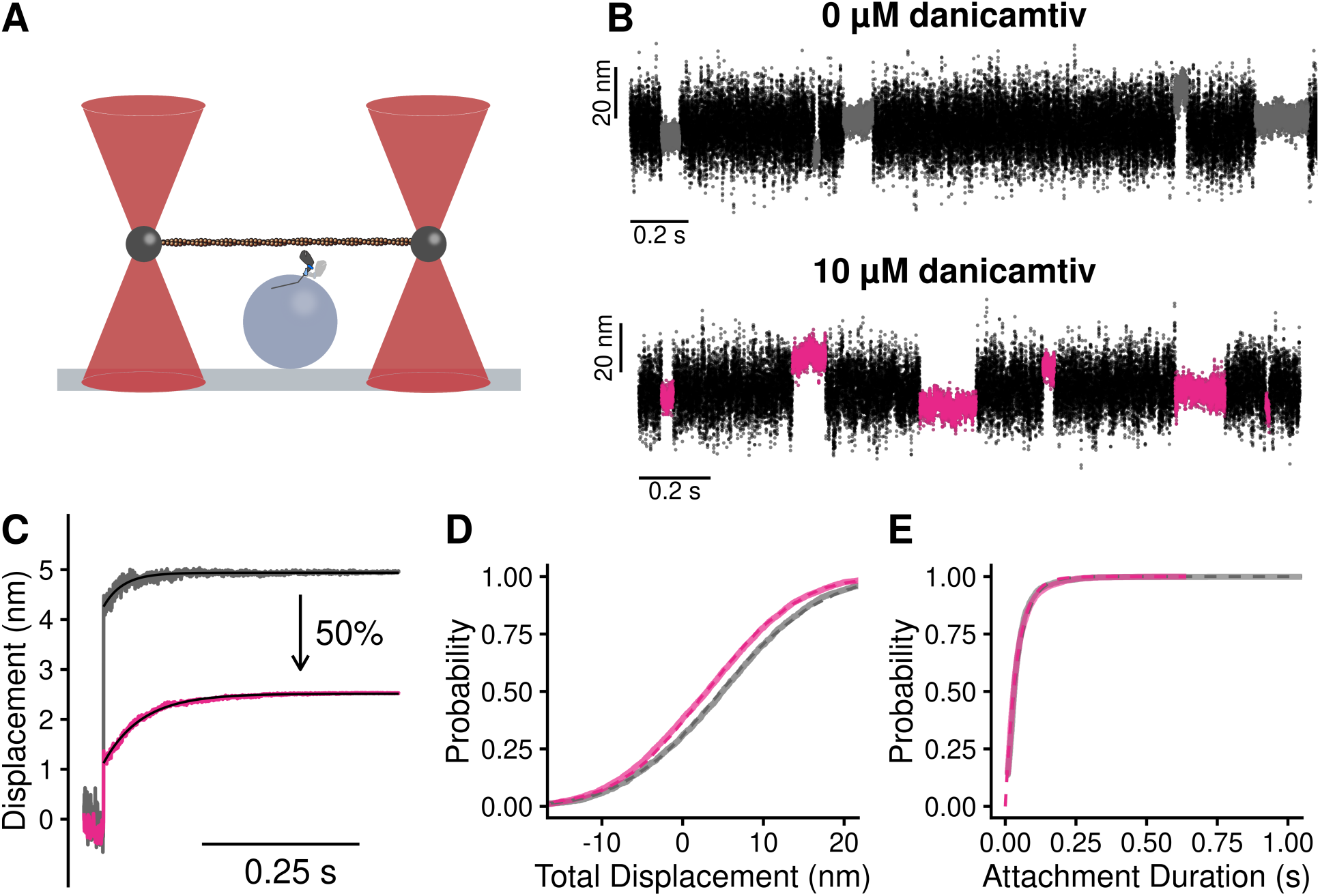
Single molecule optical trapping reveals that danicamtiv reduces the size of myosin’s working stroke without altering detachment kinetics. Black = DMSO control. Pink = 10 µM danicamtiv. **A)** Cartoon schematic of the optical trapping assay. An actin filament is strung between two optically-trapped beads and lowered onto a pedestal bead sparsely bound with myosin. **B)** Optical trapping data traces showing the stochastic binding of myosin to actin. Binding interactions are shown in grey or pink and detached states are shown in black. **C)** Time forward ensemble averages of myosin’s working stroke reveal a ∼50% reduction in the size of myosin’s total working stroke in the presence of danicamtiv. **D)** The cumulative distribution of the total working stroke displacements at 10 µM ATP is well fit by a single cumulative Gaussian function (dotted lines) with average values of 4.9 ± 9.7 nm versus 3.0 ± 9.0 nm for DMSO control and 10 µM danicamtiv, respectively (P < 0.001 using a two-tailed T-test). N = 2076 binding interactions for control and 4776 binding events for 10 µM danicamtiv. **E)** The cumulative distributions of attachment durations at 10 µM ATP. Single exponential functions were fit to the distributions using maximum likelihood estimation. 95% confidence intervals were calculated using bootstrapping methods. There is no statistical difference between control and 10 µM danicamtiv, 23 (−2.5/+2.5) s^-1^ vs. 24 (−0.8/+0.9) s^-1^ (P = 0.48). For all trapping values, see **Table 2**.

**Table 2.**
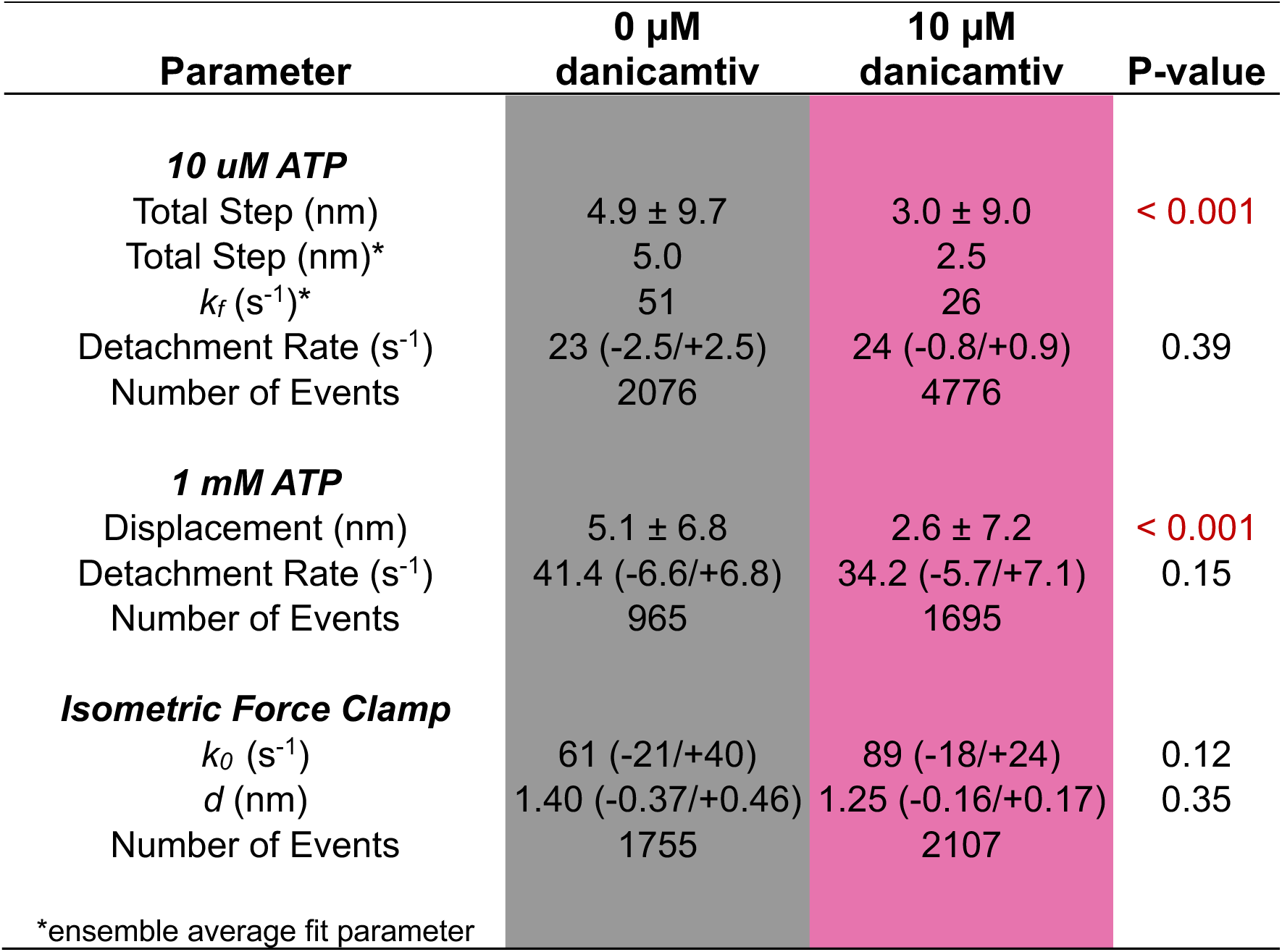
Optical trap summary.

First, we measured single molecule interactions between actin and myosin at low, non-physiological ATP concentrations (10 µM ATP) to facilitate the observation of substeps of the myosin working stroke. The size of the working stroke can be measured by fitting single exponential functions to the time forward ensemble averages (**Fig. 3C**) or by calculating the relative position change that occurs with each actomyosin interaction (**Fig. 3D**). Both analyses provide similar results (**Table 2**). We found that in the absence of danicamtiv, cardiac myosin has a working stroke size of 5.0 nm, consistent with previous measurements of human and porcine cardiac myosins (15–17, 26). We found that in the presence of danicamtiv, the total size of myosin’s working stroke was reduced to 2.5 nm (P < 0.01; **Fig. 3C**). This ∼50% reduction in the working stroke displacement is consistent with the 55% decrease in speed we observed in motility (**Fig. 1B**).

Optical trapping also enables the direct measurement of the actomyosin attachment duration. Cumulative distributions of attachment durations were generated and fitted with a single exponential function to obtain the rate of actomyosin detachment (**Fig. 3E**). We found that the detachment rate in the absence of danicamtiv at 10 µM ATP was 23 (−2.5/+2.5) s^-1^, consistent with the expected rate of detachment based on the stopped flow measurements (**Fig. 2**). Moreover, we found that there was no statistically significant difference in the actomyosin detachment rate in the presence of danicamtiv at 10 µM ATP compared to the DMSO control (24 (−0.8/+0.9) s^-1^, P = 0.39, **Fig. 3E**), consistent with our stopped flow measurements which suggested that danicamtiv does not change actomyosin detachment kinetics. This result contrasts with OM. Consistent with previous studies, we show that OM further reduces the size of the working stroke compared to danicamtiv and significantly slows the rate of actomyosin dissociation in the optical trap (**Supp.** Fig. 1) (15). Taken together, our results suggest that unlike OM, danicamtiv does not affect the kinetics of actomyosin detachment at the single molecule level at low ATP.

To ensure that our observations in the optical trap at 10 µM ATP were not a result of working at low ATP concentrations, we also collected an additional optical trapping dataset at a physiologically-relevant saturating ATP concentration (1 mM) (**Supp.** Fig. 2). At 1 mM ATP we observed that myosin’s displacement is reduced ∼50% with danicamtiv (P < 0.01). Moreover, we do not observe a difference in the actomyosin detachment rate with danicamtiv (P = 0.15, **Table 2**), consistent with our observations at low ATP and the stopped flow measurements.

Next, we tested whether danicamtiv affects myosin’s load-dependent detachment kinetics at the level of single molecules at 1 mM ATP (**Fig. 4A**). To do this, we used a feedback loop to exert force on the myosin during its working stroke, and we measured the actomyosin attachment duration under load (16). The relationship between force and attachment duration can be fit to extract the rate of the primary force-sensitive transition in the absence of force, k_0_, and the distance to the transition state (a measure of force sensitivity) (**Fig. 4B**) (27). We found that at saturating ATP concentrations, the rate of the primary force sensitive transition in the absence of danicamtiv was 61 (−21/+40) s^-1^, consistent with the rate of ADP release measured in the stopped flow (**Fig. 2B**) and previous measurements (16, 26), and this rate was not statistically different in the presence of danicamtiv (89 (−18/+24) s^-1^, P = 0.12). Moreover, the distance to the transition state was not statistically different in the absence or presence of danicamtiv (d = 1.4 (−0.37/+0.46) nm vs 1.25 (0-.16/+0.17) nm, respectively, P = 0.35). Thus, danicamtiv does not change the load-dependent kinetics of cardiac myosin at physiologically-relevant ATP concentrations, and the observed reduction in motile speed (**Fig. 1B**) can be explained by a reduction in the size of the myosin working stroke (**Fig. 3C**).

**Figure 4.**
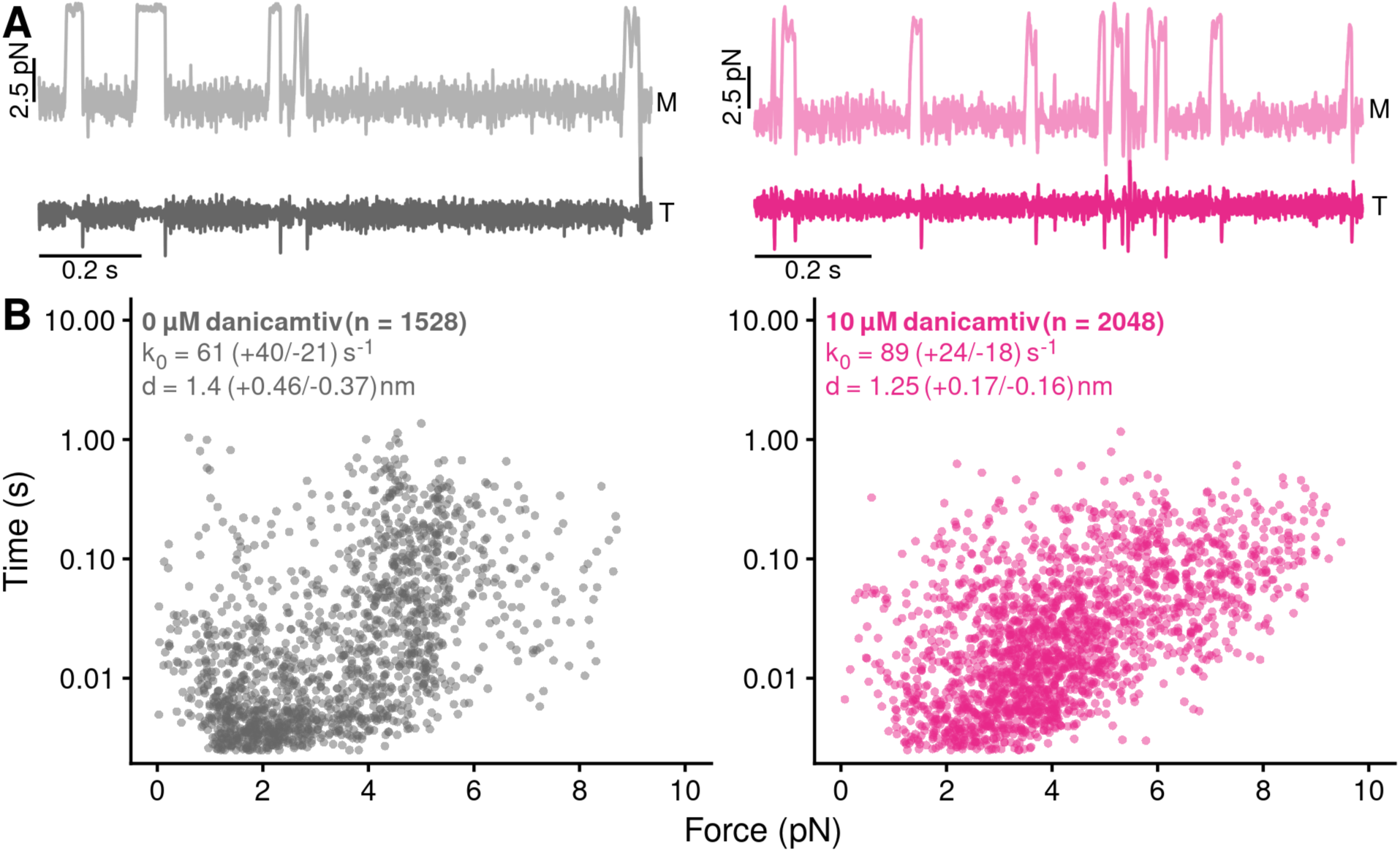
Danicamtiv does not alter myosin’s load-dependent detachment kinetics at 1 mM ATP. Black = DMSO control. Pink = 10 µM danicamtiv. **A)** An isometric force clamp was used to maintain actin at an isometric position during myosin binding interactions. To do this, the motor bead (M) was moved to hold the transducer bead (T) at an isometric position. Data traces are shown. **B)** Plots of actomyosin attachment duration versus the average resistive force applied during the binding event. Data are exponentially distributed at each force. Each point represents an independent actomyosin binding interaction. The data were fitted with the Bell equation using maximum likelihood estimation and 95% confidence intervals were calculated for each parameter by bootstrapping. The detachment rate in the absence of load, *k_0_,* was not different between control and 10 µM danicamtiv, 61 (−21/+40) vs 89 (−18/+24) s^-1^ (P = 0.17). These values are consistent with our measurements of the rates of ADP release from stopped-flow experiments. The distance to the transition state, *d*, which measures the load-sensitivity of the detachment rate, was not different between control and 10 µM danicamtiv, 1.40(−0.37/+0.46) vs 1.25 (−0.16/+0.17) nm (P = 0.43).

### Danicamtiv increases the kinetics of actomyosin attachment

Our results clearly demonstrate that at the level of single crossbridges, danicamtiv does not affect actomyosin detachment kinetics. Therefore, we investigated whether danicamtiv affects actomyosin attachment kinetics. Our steady-state ATPase measurements (**Fig. 1A**) demonstrate that danicamtiv increases the overall steady-state ATPase rate, indicative of the fact that danicamtiv increases the rate of the slowest step of the ATPase cycle. For β-cardiac myosin cycling in the presence of actin, the overall cycle rate is limited by attachment kinetics (23). Since danicamtiv increases the steady state ATPase without altering detachment kinetics, we posited the increase in ATPase could be a result of danicamtiv accelerating attachment kinetics.

To test whether danicamtiv affects attachment kinetics, we used stopped flow techniques to measure the rates of actomyosin attachment and detachment under single turnover conditions (28) (**Fig. 5A**, see Supplemental Materials for additional details). Myosin was preincubated with an excess of pyrene-labeled actin to form a rigor complex that quenches pyrene’s fluorescence. The actomyosin was then rapidly mixed with a sub-saturating concentration of ATP, causing an increase in fluorescence that reports the actomyosin detachment rate at this ATP concentration, *k_det_*. A low concentration of ATP is used to ensure that this process only occurs once (i.e., a single turnover). Once off actin, myosin hydrolyzes ATP and then re-attaches to actin, quenching the pyrene fluorescence and reporting the rate of myosin attachment at this actin concentration, *k_att_*.

**Figure 5.**
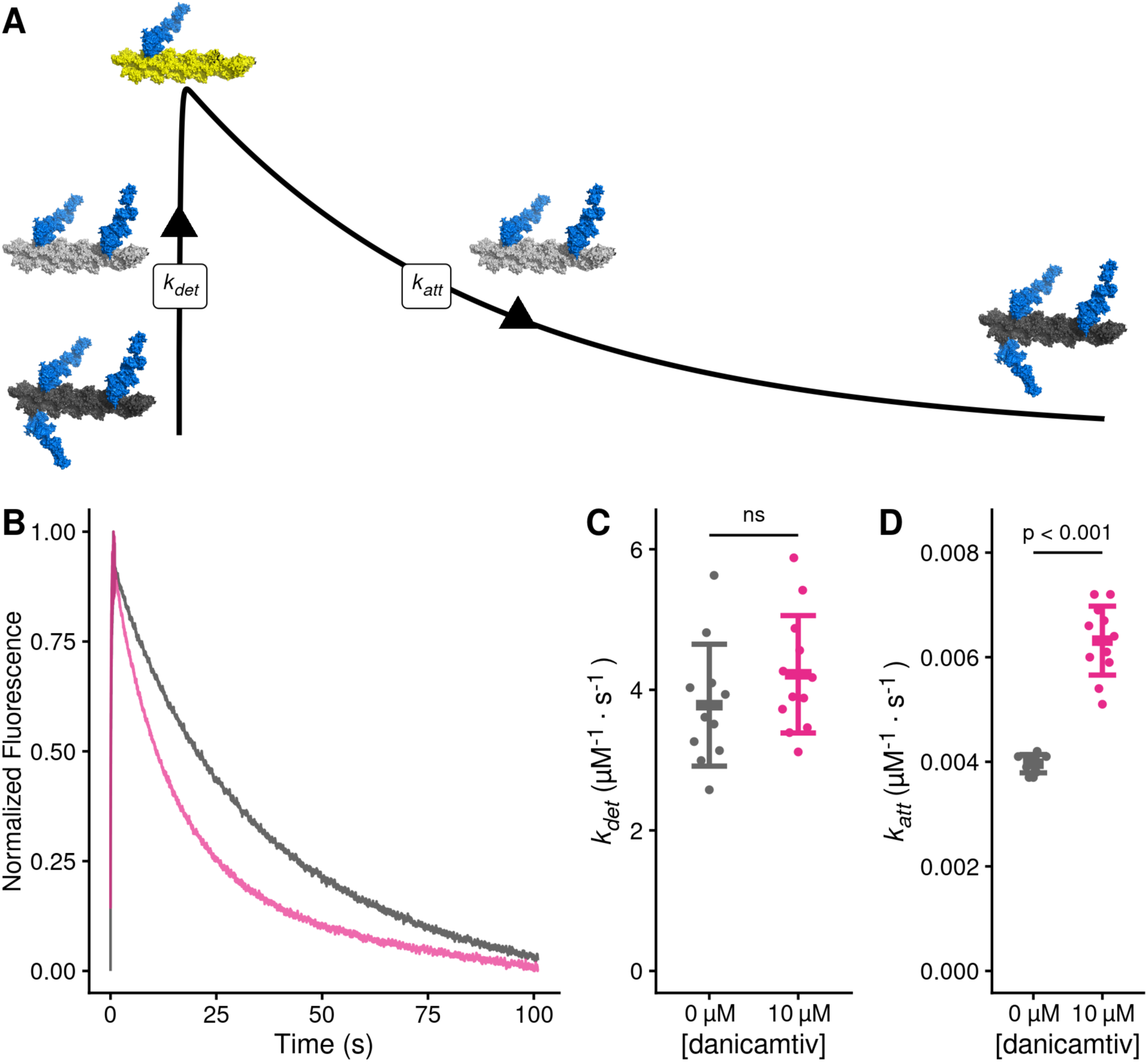
Danicamtiv increases myosin’s attachment rate in a single turnover stopped flow assay. Black = DMSO control. Pink = 10 µM danicamtiv. **A)** Schematic conceptually describing the single turnover assay used to measure myosin’s attachment and detachment rates. In this assay, myosin and pyrene labeled actin are pre-incubated and then mixed with a sub-saturating concentration of ATP. The pyrene fluorescence increases as myosins detach from actin, and the increase in fluorescence reports the rate of detachment of myosin from actin (*k_det_*). Myosin then reattaches to actin, quenching the fluorescence and reporting the attachment rate (*k_att_*). **B)** Fluorescence transients from the single turnover assay. Data were fitted as described in the Supplemental Methods. **C)** The average second-order rate of detachment (*k_det_*) was similar with and without 10 µM danicamtiv (3.8 ± 0.9 vs. 4.2 ± 0.8 s^-1^; P = 0.23). **D)** The second-order rate of attachment (*k_att_*) increased with the addition of 10 µM danicamtiv (0.0040 ± 0.0002 vs 0.0063 ± 0.0007 µM^-1^⋅s^-1^; P = < 0.001). For **C** and **D**, the thick lines show the average values, and the error bars show the standard deviation. The individual points are the fitted second-order rates to individual transients collected across three experimental replicates. Statistical testing was done using a two-tailed T-test after passing a normality test.

The single turnover fluorescence transients consisted of two phases, where the rate of detachment was faster than the rate of attachment, consistent with the notion that the rate of attachment limits the overall ATPase cycle time (**Fig. 5B, Supp. Figs. 3 and 4**). We saw that there was no statistically significant difference in the rate of detachment in DMSO or danicamtiv. (3.8 ± 0.9 vs 4.2 ± 0.8 s^-1^, respectively, P = 0.23, **Fig. 5C**). This measured detachment rate is in agreement with the second-order rate of ATP-induced dissociation at 0.75 µM ATP measured in the stopped flow (**Table 1**). We also saw that the observed attachment rate was significantly faster with 10 µM compared to DMSO controls (0.0040 ± 0.0002 vs 0.0063 ± 0.0006 µM^-1^⋅s^-1^, respectively, P < 0.001, **Fig. 5D**). Taken together, our data demonstrate that danicamtiv increases attachment kinetics without affecting detachment kinetics.

### Danicamtiv-induced increase in the myosin attachment rate causes increased thin filament activation

Given that danicamtiv increases the attachment rate of myosin to actin, we hypothesized that this would lead to an increase in the fraction of myosin heads bound to actin (i.e., the duty ratio). To test this hypothesis, we used a well-established variation of the *in vitro* motility assay where the speed of actin filament movement was measured as a function of surface myosin (29, 30). Actin filaments move at their maximal speed if there is a sufficient concentration of myosin on the surface to ensure that at least one myosin head is attached to the filament at any given time. We observed that despite moving slower, danicamtiv-treated myosin required less myosin to reach saturation, consistent with an increased duty ratio with drug (**Figs. 6A and B**).

**Figure 6.**
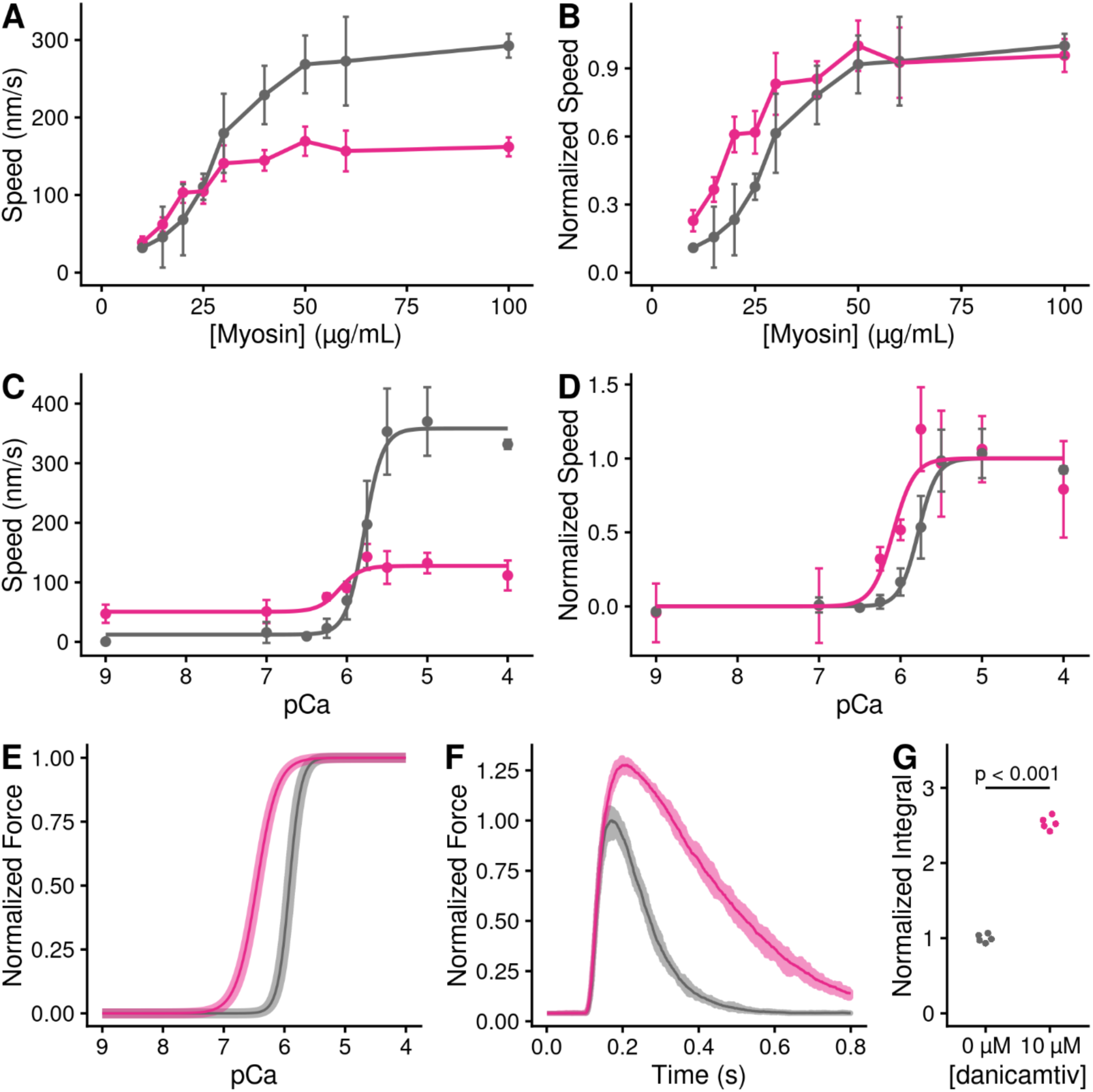
Danicamtiv increases motility speed in the presence of regulatory proteins and these effects on muscle contraction can be recapitulated computationally. **Black** = DMSO control. Pink = 10 µM danicamtiv. **A and B)** Unregulated *in vitro* motility speed as a function of the myosin concentration on the flowcell surface. The speed decreases if there is not at least one active myosin head bound to actin at any given time. Thus, if the duty ratio increases with drug, less myosin will be required to reach saturation. 10 µM danicamtiv decreases motility speed at higher myosin concentrations but increases speed at low myosin concentrations, indicative of a higher duty ratio, despite having a smaller working stroke. **A)** shows the measured speed and **B)** shows normalized data. ∼40 filaments were tracked across four fields of view from two different experimental preparations. **C and D)** Regulated i*n vitro* motility speed using thin filaments decorated with troponin and tropomyosin as a function of calcium. The data were fitted with the Hill equation and the fitted values ± standard error are: V_max_ values are 386 ± 8 vs. 135 ± 7 nm/s (P < 0.001), pCa50 values are 5.76 ± 0.02 vs. 6.1 ± 0.09 (P = 0.01), and the Hill coefficients are 3.4 ± 0.4 vs 3.1 ± 1.5 (P = 0.85) for the control vs. 10 µM danicamtiv, respectively **C)** Shows the measured speed and **D)** shows the data normalized to the fitted V_max_ and V_min_. Each point represents average speed with error bars showing the standard deviation of ∼40-60 filaments imaged from 4-6 fields of view from 2-3 experimental replicates. **E)** Simulated force-calcium relationship from FiberSim. To simulate danicamtiv, we incorporated increased actin attachment, reduced myosin working stroke, and an increase in the population of active myosin heads. The simulations recapitulate the shift seen in the motility experiments. 5 replicates were conducted, the shaded region shows the range of values, and the solid line shows the mean. **F)** Simulated twitch in response to a calcium transient using the same simulation parameters. Danicamtiv increases the maximal force, slows kinetics, and **G)** increases the force-time integral.

Given the increase in the rate of myosin binding to actin with danicamtiv, we hypothesized that danicamtiv would increase myosin-induced thin filament activation, since thin filament activation depends on myosin binding (31). To test this, we reconstituted thin-filaments in the *in vitro* motility assay, and we measured the speed of myosin-based translocation over a range of calcium concentrations. We saw that danicamtiv increased thin filament motility at submaximal calcium levels where the motility speed is limited by the rate of myosin attachment (e.g. at pCa 7, 16 ± 28 vs. 51 ± 31 nm/s, P < 0.001) (**Fig. 6C**). These low calcium levels are within the physiological range of calcium concentrations seen in muscle (32). Taken together, our data suggest that danicamtiv-induced increases in attachment kinetics lead to increased thin filament activation at submaximal calcium concentrations.

### Computational modeling connects molecular effects and muscle function

To better understand how changes at the molecular scale with danicamtiv would translate into altered muscle contraction, we used computational modeling. We used a spatially explicit model of muscle contraction, FiberSim (33). This model has previously been used to successfully model several physiologically important parameters, including the force-calcium relationship and the force generated in response to a calcium transient.

We used FiberSim to model the effects of danicamtiv on key muscle parameters based on our molecular measurements. Our results show that danicamtiv reduces the size of the working stroke while increasing the rate of crossbridge attachment. Moreover, previous X-ray diffraction studies have shown that danicamtiv causes myosin to transition from an autoinhibited interacting heads motif to an activated disordered relaxed state (14). We adjusted the model input parameters to match these changes, and we simulated the effects of each of these changes in isolation (**Supp.** Fig. 5) and all together (**Figs. 6E and F**). We simulated both the force-calcium relationship and the force generated in response to a calcium transient. When looking at the composite effects of all three changes, we see that this causes a shift in the force calcium relationship towards submaximal calcium activation (**Fig. 6E**) that agrees well with our experimental measurements (**Fig. 6D**). Moreover, the modeling shows that danicamtiv is expected to increase both the magnitude of the force generated in response to a calcium transient with slight slowing of both the rates of force development and relaxation (**Fig. 6F**). Finally, the modeling predicts that danicamtiv will increase the force-time integral (**Fig. 6G**). Taken together, our modeling provides insights into how danicamtiv affects the kinetics and mechanics of myosin, leading to increased muscle function.

## DISCUSSION

### At the level of single crossbridges, danicamtiv’s biophysical mechanism is more complicated than a “myosin activator”

Our observation of ∼50% reduced motility with danicamtiv (**Fig. 1B**) is consistent with the excellent study by Kooiker et al. (14). Previously, it was suggested that this reduced speed could be due to danicamtiv’s effects on ADP binding and release, since the rate of ADP release limits the shortening speed of muscle (34). We directly measured the rate of ADP binding, rate of ADP release, and the equilibrium constant for ADP binding, and we do not see changes in any of these parameters (**Figs. 2B and C**). Moreover, we directly measured the rate of actomyosin dissociation in the optical trap at both high and low ATP concentrations, and we did not observe any changes in the detachment rate (**Fig. 3E**). We also measured the load-dependent detachment kinetics in the optical trap, and we did not observe any changes with danicamtiv treatment (**Fig. 4**). Finally, we used a single turnover stopped flow assay, and we do not see any difference in the detachment kinetics (**Fig. 5C**) Taken together, our results demonstrate that danicamtiv does not have effects on the rate of ADP release or actomyosin detachment at the level of single crossbridges.

The speed in the motility assay is proportional to the step size of the myosin divided by the amount of time that the crossbridge remains attached (19). While we did not observe any changes in attachment time in the optical trap, we did observe a ∼50% decrease in the size of the myosin working stroke (**Fig. 3E**). As such, the ∼50% decrease in motility speed can be explained by a ∼50% reduction in the size of the working stroke. Taken together, at the level of individual crossbridges, the reduced speed in motility seen with danicamtiv cannot be explained by changes in detachment kinetics, rather, it is due to a decrease in the size of the myosin working stroke.

Danicamtiv was initially discovered in a high-throughput screen for molecules that activate the steady-state ATPase activity of myosin, and as such, it was initially classified as a myosin activator (11); however, as we show here, it has a more complex biophysical mechanism at the level of single crossbridges. The ATPase assay uses a minimal number of components (i.e., myosin, actin, and ATP) to measure the steady-state rate of ATP turnover. While this assay is useful for drug screening due to its well-characterized and easily measured outputs, it also has important limitations due to its simplified nature. This assay considers only the effects of drugs on kinetics, and it does not consider effects on mechanics or incorporate higher-order structures that are important for muscle function (e.g., sarcomere lattice, myosin autoinhibition in the thick filament, calcium-based regulation). This limitation becomes clear in the case of danicamtiv, where mechanics and kinetics are uncoupled. We show that danicamtiv is an activator of myosin’s steady-state ATPase rate (**Fig. 1A**), but an inhibitor of myosin mechanics (**Fig. 3C**). This demonstrates that danicamtiv partially uncouples myosin mechanics and kinetics, and its biophysical mechanism is more complicated than a simple myosin activator. We propose that danicamtiv should be classified as a myosin-binding sarcomeric activator.

### Danicamtiv’s activating properties emerge in higher-order structures

Since danicamtiv has both inhibitory and activating effects at the level of individual crossbridges, we further investigated its effects in higher-order structures. Increased cardiac contractility has been observed in muscle fibers, small animal models, and humans (9, 11–14, 35). One difference between actomyosin in isolation and in muscle is the presence of the thin filament regulatory proteins, tropomyosin and troponin. In the absence of calcium, tropomyosin blocks the myosin strong binding sites on actin, preventing myosin attachment and subsequent force generation (36). During systole, calcium binds to troponin, leading to movement of tropomyosin followed by attachment of myosin crossbridges. Thus, the process of thin filament activation depends both on both calcium and myosin binding to the thin filament.

Here, we observed that danicamtiv increases the rate of myosin attachment to actin (**Figs. 1A and 5D**). As such, we hypothesized that increased attachment would lead to increased binding to the thin filament, causing activation of the thin filament at submaximal calcium levels. In fact, this is what we observe in the *in vitro* motility assay using regulated thin filaments, and similar shifts were seen in muscle fiber experiments (12–14). To test whether this increased myosin attachment could contribute to the observed changes in muscle fiber force, we performed computational modeling of the sarcomere using FiberSim (**Fig. 6E**). We found that changing myosin’s rate of attachment to the thin filament alone is sufficient to recapitulate the shift towards submaximal calcium activation that we observed in the *in vitro* motility assay (**Supp.** Fig. 5). Taken together, we propose that danicamtiv increases muscle contraction, in part, through activation of the thin filament.

As mentioned above, simplified systems cannot capture all aspects of cardiac contraction, and previous studies in muscle fibers have demonstrated several effects that we cannot observe at the level of individual crossbridges (12–14). To gain some insights into how changes at the level of individual crossbridges translates into muscle function, we used a spatially explicit model of muscle contraction to simulate key physiological parameters (**Figs. 6E and F** and **Supp.** Fig. 5). We simulated the individual and composite effects of (1) increased crossbridge attachment based on **Figs. 1A** and **5D**, (2) decreased working stroke size based on **Figs. 1B** and **3C**, and (3) an increase in the number of myosin crossbridges available to interact with the thin filament based on X-ray diffraction studies of muscle fibers (14). Our results clearly demonstrate that these danicamtiv-induced changes at the level of single crossbridges are sufficient to reproduce the shift towards submaximal calcium activation, increased peak force production in response to a calcium transient, and an increase in the force-time integral. Moreover, we observe slightly slowed rates of force development and relaxation with danicamtiv that emerge without changes in the rate of crossbridge detachment. Taken together, our results demonstrate the importance of multiscale studies for understanding the mechanisms of small molecules targeting myosin.

### Comparison with omecamtiv mecarbil

The first compound targeting cardiac myosin for the treatment of systolic heart failure was omecamtiv mecarbil (OM) (5, 6). Like danicamtiv, OM was also identified in a screen for small molecules that increase myosin’s steady-state ATPase activity. OM made it to stage III clinical trials, but ultimately the FDA declined to approve OM due to its limited effect size and its impact on cardiac relaxation (10). In particular, patients treated with OM showed prolonged systole and impaired relaxation, which lead to an increase in serum troponin, indicative of cardiac damage.

There are some similarities between OM and danicamtiv seen at the molecular scale. Both danicamtiv and omecamtiv increase myosin’s ATPase activity and slow the rate of motility (5, 37). Both danicamtiv and OM decrease the size of the myosin working stroke (15); however, the reduction in the size of the working stroke with danicamtiv was not as severe as the reduction caused by OM (**Supp.** Fig. 1C). The decrease in size of the working stroke can be seen for both OM and danicamtiv; however, this effect is more pronounced for OM which almost completely eliminates the working stroke.

The key difference between OM and danicamtiv is that omecamtiv significantly increases the amount of time that actin and myosin remain attached while danicamtiv does not. This can be seen in **Supp.** Fig 1D. This subtle difference has important implications for the mechanism of action. OM increases the attachment duration, leading to slowed detachment of myosins, prolonging systole. This effect would not be expected for danicamtiv, suggesting that it might have better effects on relaxation and diastolic function. Consistent with this notion, experiments using engineered heart tissues and muscle fibers showed that danicamtiv has a lower impact on relaxation than OM (9, 13). It is worth noting however, that while the effects of danicamtiv on diastolic function are smaller that OM, they still exist (9, 35), and we can observe evidence of slowed relaxation in our computational modeling. It remains a challenge to the field to develop small molecules that can uncouple myosin’s effects on systole and diastole.

### Conclusions

Here, we demonstrate the effects of danicamtiv on single crossbridges and highlight how properties at the single molecule level translate into effects seen in muscle fibers. Importantly, our results demonstrate that danicamtiv is a myosin binding sarcomeric activator that exerts its effects in part through thin filament activation.

## Acknowledgements

The authors acknowledge financial support provided by the National Institutes of Health (R01 HL141086 to M.J.G., R01 HL148785 to K.S.C., T32 HL125241 to B.S.), the Children’s Discovery Institute of Washington University and St. Louis Children’s Hospital (PM-LI-2019-829 M.J.G.), and the American Heart Association (TPA 970198 to M.J.G).

## Conflict of interest statement

All experiments were conducted in the absence of any commercial or financial relationships that could be construed as potential conflicts of interest. M.J.G. discloses research funding from Edgewise Therapeutics on an unrelated project.

## Author contributions

Conception and oversight by M.J.G.. ATPase experiments were conducted by L.G.. Computational modeling was conducted by C.S., K.S.C., and B.S.. All single molecule and transient kinetic experiments were conducted by B.S.. All authors contributed to the analysis of the data. The first draft was written by M.J.G.. All authors contributed to the writing and/or editing of the manuscript.

## MATERIALS AND METHODS

Full experimental procedures can be found in the Supplemental Materials.

### Biochemical Kinetic Measurements

Porcine ventricular actin and myosin were purified from tissue, and human troponin and tropomyosin were recombinantly expressed in *E. coli* (22). Actin was labeled with pyrene as previously described (22). Danicamtiv was purchased from Selleckchem (99.1% purity, S9948). ATPase measurements as a function of actin concentration were conducted using the NADH enzyme coupled assay (21, 38). Stopped flow measurements were conducted in an SX-20 instrument (Applied Photophysics). Using these techniques, we measured the rates of ATP induced actomyosin dissociation, ADP release, ADP hydrolysis, ADP binding affinity, and single turnover kinetics (21).

### *In Vitro* Motility and Optical Trapping Experiments

*In vitro* motility assays were conducted as previously described, where actin filaments translocate over a bed of myosin in the presence of ATP (39, 40). Thin filament regulation was reconstituted by adding calcium and the regulatory proteins troponin and tropomyosin. Optical trapping was done using the three-bead assay in which an actin filament is stretched between two optically trapped beads and lowered on to a pedestal bead that is sparsely coated with myosin (16, 24, 27). Data were analyzed using the SPASM software (24).

### Computational modeling

Computational modeling was done using FiberSim, a spatially explicit model of muscle contraction (33). Model input parameters were modified to match experimental measurements. The force-calcium relationship and the time-dependent response to a calcium transient were simulated.

### Data Availability and Reproducibility

All data is included in the project repository hosted on Zenodo (<u>10.5281/zenodo.12636349</u>). All analysis was performed in R version 4.4.0 (R Core Team) unless otherwise noted. The code has been made available to reproduce verbatim all figures and related analyses in the project repository.

## Notes

### Competing Interest Statement

The authors have declared no competing interest.

### Summary of Updates

This manuscript includes additional experimental replicates compared to the original document.

