## Supplemental Materials for "Danicamtiv reduces myosin’s working stroke but enhances contraction by activating the thin filament"

### Supplemental Methods

#### Proteins and Solutions

Full length β-cardiac myosin and actin were purified from porcine left ventricular tissue as we have previously done (1). The proteolytic cleavage of full-length myosin into myosin sub-fragment 1 (S1) by chymotrypsin was performed as described by previously described (2, 3). Actin was prepared from acetone powder as previously described (4). Actin was labeled with n-(1-pyrene)iodoacetamide for stopped flow experiments as previously described (5). All actin was phalloidin stabilized in a 1.1:1 ratio. Human troponin and tropomyosin were expressed in E. coli and purified as previously described (2). All experiments were performed in KMg25 buffer unless otherwise noted (25 mM KCl, 10 mM EGTA, 60 mM MOPS pH 7.0, 1 mM DTT, and 5 mM $MgCl_2$). Danicamtiv was purchased from Selleckchem and dissolved in DMSO (99.1% purity, S9948) and diluted in KMg25 to a final concentration of 10 μM in the final assay buffers. The experiments were conducted with 0.1% DMSO controls.

#### Steady-State ATPase Measurements

The steady-state ATPase rate of myosin was measured using the NADH-linked assay as we have previously done (6-8). The NADH-linked ATPase measurements were conducted with myosin S1 in a specific ATPase buffer containing 20 mM Imidazole pH 7.5, 10 mM KCl, 2 mM $MgCl_2$, and 1 mM DTT assayed in a BioTek Synergy H1M plate reader using a 96 well plate with shaking at 25°C. Absorbance was monitored at 340 nm continuously over 10 minutes. The contribution of actin alone to the ATPase rate was subtracted. Data showing the rate as a function of actin concentration were fitted using the Michaelis-Menten equation.

**Stopped Flow Transient Kinetics**

Stopped flow measurements were conducted in an SX-20 instrument from Applied Photophysics. All nucleotide concentrations were determined spectroscopically. All stopped flow assays used myosin S1. Experiments were conducted in KMg25 at 20°C. For the measurements of ATP-induced dissociation, ADP release, ADP affinity, and single turnovers, pyrene-labeled actin was used with an excitation wavelength of 365 nm and a 395 nm filter before the photomultiplier tube. Measurement of ATP hydrolysis was done using intrinsic tryptophan fluorescence, with an excitation wavelength of 295 nm and a 320 nm filter placed before the photomultiplier tube. All concentrations are given before mixing.

We measured the rate of ADP release from actomyosin $k_{+5}$ as we have previously done (1, 2, 5, 9). Briefly, 1 μM myosin, 1 μM pyrene actin, and 100 μM Mg.ADP were pre-incubated in syringe 1 and rapidly mixed 5000 μM Mg.ATP. Resultant fluorescence transients were well fitted by a single exponential function to provide the rate of ADP release.

To measure the ADP affinity, $K_5$', we pre-incubated 1 μM myosin, 1 μM pyrene actin, and a variable amount of Mg.ADP. We then rapidly mixed this with 100 μM Mg.ATP. Fluorescence transients were well fitted by single exponential functions. The observed rate was plotted as a function of ADP, and the resultant curve was well fitted by a hyperbolic curve:

$$k_{obs} = \frac{k_0}{1 + \frac{[ADP]}{K_5{}'}}$$

where $k_0$ is the rate in the absence of ADP and $K_5$ is the ADP affinity. From the measured rate of ADP release and the ADP affinity, it was possible to calculate the rate of ATP binding to actomyosin, $k_{-5}$.

The rate of ATP-induced actomyosin dissociation was measured as we have previously done (1, 2, 5, 9). 1 μM myosin, 1 μM pyrene actin, and 0.04 U/mL apyrase VII were pre-incubated in syringe 1 and then rapidly mixed varying concentrations of Mg.ATP. Fluorescence transients were well fitted by the sum of two exponential functions, as previously described (6). The fast phase reports the rate of ATP binding and subsequent actomyosin dissociation. The slow phase reports on a well-established nucleotide isomerization. The amplitude of the fast phase was fixed at low ATP concentrations because some signal is lost in the dead time at higher concentrations of ATP. The observed rate of the fast phase increases hyperbolically with ATP concentration, and the data was fitted with the hyperbolic function:

$$k_{fast} = \frac{k_{+2}{}'[ATP]}{K_1{}' + [ATP]}$$

The observed rate of the slow phase at saturating ATP concentrations reports the rate of the nucleotide isomerization $k_{+\alpha}$. The ratio of the fast and slow amplitudes can be used to calculate the equilibrium constant for the nucleotide free isomerization, $K_\alpha$. The reverse rate constant can be calculated from the forward rate and the equilibrium constant.

To measure the rate of ATP binding and hydrolysis to myosin, we used the intrinsic tryptophan fluorescence of myosin that changes with hydrolysis (6). We preincubated 2 μM myosin with 0.04 U/mL apyrase VII and then rapidly mixed this with a range of ATP concentrations. The individual fluorescence transients were well fitted by single exponential functions, and the relationship between the observed rate and the ATP concentration was well fitted by a hyperbolic function.

**Single Turnover Measurements of Actomyosin Detachment and Attachment**

To measure the rates of actomyosin detachment ($k_{det}$) and attachment at a given concentration of actin ($k_{att}$), we used a single turnover measurement (10). For practical experimental reasons, this experiment could not be performed under optimal pseudo-first order conditions. As such, the data show some deviation from single exponential behavior, and it is more appropriate to solve the system of differential equations. We mixed 2.5 μM myosin and 10 μM pyrene labeled actin (all concentrations after mixing) to form 2.5 μM actomyosin (AM) and 7.5 μM free pyrene actin (A** where the stars denote fluorescence). We monitored the change in pyrene fluorescence since free pyrene actin has high fluorescence while myosin bound to pyrene actin has quenched fluorescence. We then rapidly added 0.75 μM ATP (T). ATP binding to actomyosin causes rapid detachment of the myosin, which can be seen by an increase in fluorescence. Then ATP bound to myosin undergoes rapid hydrolysis to myosin*ADP*Pi (MDP). MDP can then reattach to the actin, quenching the fluorescence and releasing phosphate (P).

Data were analyzed with the following overall kinetic scheme:

$$AM + T + A^{**} \xrightarrow{k_{det}} MDP + AM + A^{**} \xrightarrow{k_{att}} AM + P + A^{**}$$

The reaction followed the set of differential equations:

$$\frac{d[AM]}{dt} = -k_{det}[AM] * [T] + k_{att}[A^{**}] * [MDP]$$

$$\frac{d[A^{**}]}{dt} = k_{det}[AM] * [T] - k_{att}[A^{**}] * [MDP]$$

$$\frac{d[T]}{dt} = -k_{det}[AM] * [T]$$

$$\frac{d[MDP]}{dt} = k_{det}[AM] * [T] - k_{att}[A^{**}] * [MDP]$$

$$\frac{d[P]}{dt} = k_{att}[A^{**}] * [MDP]$$

The initial starting conditions were [AM] = 2.5 μM, [T] = 0.75 μM, [A] =7.5 μM, [MDP] = 0, and [P] = 0. In the stopped flow, we observe the change in the fluorescence as MDP is formed and then disappears, so our measured fluorescence signal is proportional to the [MDP]. The normalization constant of this signal was a fitting parameter.

These experiments could not be performed under pseudo-first order conditions, and therefore data needed to be analyzed by solving a set of differential equations. We used the ode45 function in Matlab and least squares optimization to solve the system of differential equations for the values of $k_{att}$ and $k_{det}$ that best fit the experimental data (as well as the normalization constant to convert fluorescence signal to concentration) (**Supplemental Figures 3 and 4)**.

Consistent with our transient kinetic and optical trapping experiments, the single turnover transients demonstrate that 10 μM danicamtiv does not change detachment kinetics (**Supplemental Fig. 4C**). Moreover, these experiments demonstrate that 10 μM danicamtiv increases the attachment kinetics (**Supplemental Fig. 4D**). It is important to note that these measurements could not be conducted at saturating concentrations of actin due to experimental limitations. As such, the rate of attachment here does not represent the maximal rate of attachment; however, our transient and steady-state kinetic experiments demonstrate that the overall ATPase cycle is not limited by detachment kinetics. The steady-state ATPase experiments clearly demonstrate that danicamtiv increases the ATPase rate, likely through effects on attachment kinetics.

#### *In vitro* Motility Assays

*In vitro* motility assays were performed as previously described (1, 2, 5, 9). For regulated motility, thin-filaments were reconstituted by the addition of 0.5 µM troponin and 0.5 µM tropomyosin. For the regulated motility experiments, the levels of free calcium were set using MaxChelator (11). Videos were recorded at room temperature and analyzed using MTrackJ (12) in Fiji (13).

#### Optical Trapping

All optical trapping assays were conducted on a custom-built microscope free setup as described previously (1, 14, 15). Full length myosin underwent a deadhead spin down on the day of experimentation, and it was diluted to achieve single molecule conditions (1-3 nM). The final assay buffer was KMg25 with 1 mg/mL BSA, the desired ATP concentration, 5-10 nM TRITC biotin F-actin, 1 mg/mL glucose, 192 U/mL glucose oxidase, and 48 µg/mL catalase. Experiments were performed at room temperature.

The analysis of optical trapping data was performed using automated event detection based on changes in the covariance between the optically trapped beads (14). For the 1 mM ATP trapping dataset, analysis was done using a separate software program (16) that is available as an R package and is available on GitHub (https://github.com/brentscott93/lasertrapr). At 1 mM ATP, myosin's rigor lifetime is very short lived (~1 ms) making it impossible to resolve substeps of the myosin working stroke. For the 1 mM ATP dataset, the total displacement is measured from the average position across the duration of the binding event. For all datasets, the cumulative distributions were using with maximum likelihood estimation as described previously (17).

For the load-dependent measurements of actomyosin interactions, experiments were conducted at saturating (1 mM) ATP. Data were analyzed by applying a Hidden Markov model and changepoint analysis to the position of the motor bead. The average force was calculated across the duration of the binding event. Data were fitted to the Bell equation using maximum likelihood estimation and confidence intervals were calculated by bootstrapping simulations (17).

**Computational modelling with FiberSim**

To simulate the molecular effects of danicamtiv in muscle, we used FiberSim (18). FiberSim is a spatially explicit model of muscle contraction based on fundamental biophysical properties. We used a previously validated base model to serve as a reference control. To model the molecular effects of danicamtiv, we changed three parameters in the model (1) the myosin step size, (2) the population of active myosin heads, and (3) the attachment rate. We examined the effects of changing these parameters both in isolation and in combination. All the configuration and relevant model files are in the project repository hosted on Zenodo. FiberSim 2.1.0 was used to run the simulations on a Windows 10 computer.

**Statistical Analysis**

All data was collected over multiple days from at least 2 independent protein preparations. For the ATPase, stopped flow measurements, and motility experiments, parameter values for each day were calculated from fitting of the data and the reported uncertainties are from the analysis of different data sets. Normally distributed data sets were analyzed using 2-tailed student's T-tests. Data that were not normally distributed were analyzed using the non-parametric Mann-Whitney test. For the optical trapping experiments, data were analyzed using maximum likelihood estimation followed by estimation of 95% confidence intervals by 1000 rounds bootstrapping (17).

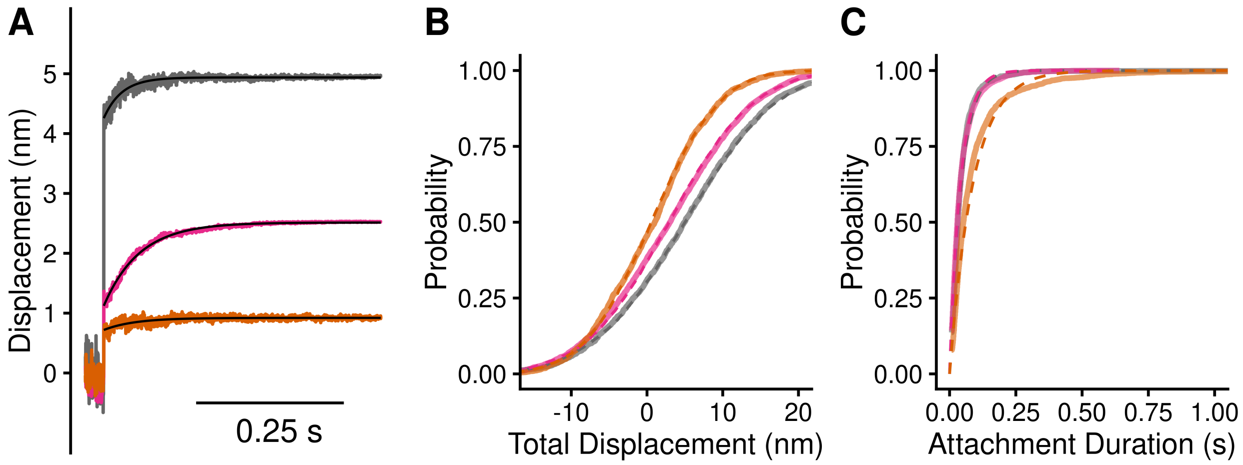


**Supplemental Figure 1: Single molecule optical trapping at 10 μM ATP with omecamtiv mecarbil (OM).** This is the same data as main text Fig. 3 with the addition of a dataset collected in the presence of 10 μM OM. The OM dataset consists of a total of 2225 single molecule binding events. Black = DMSO control. Pink = 10 μM danicamtiv. Orange = 10 μM OM. **A)** Total displacements of the time forward ensemble averages. **B)** Cumulative distributions of myosin's total working stroke size. Dotted lines are fits to a cumulative gaussian distribution. Myosin's working stroke was 4.9 ± 9.7 nm in the absence of drugs. Danicamtiv reduced the working stroke to 3.0 ± 9.0 nm. OM almost eliminated the working stroke (0.7 ± 7.1 nm). A one-way ANOVA was used to test for significance ($P < 0.001$) and Tukey post-hoc was used for pairwise comparison. All comparisons were significantly different ($P < 0.001$ for each pairwise comparison). **C)** Cumulative distribution of attachment durations. Single exponential functions were fit to the distributions using maximum likelihood estimation. 95% confidence intervals were calculated using bootstrapping methods. There is no statistical difference between control and 10 μM danicamtiv, 23 (-3/+3) $s^{-1}$ vs. 24 (-1/+1) $s^{-1}$ ($P = 0.48$). Unlike danicamtiv, OM slows the actomyosin detachment rate to 11 (-1/+1) $s^{-1}$. A Kruskal-Wallis ranks test (non-parametric) was used to compare all groups ($P < 0.001$) and Dunn's post-hoc used for pairwise comparison. The control and danicamtiv conditions were both significantly different than OM ($P < 0.001$ for those pairwise comparisons with Sidak correction).

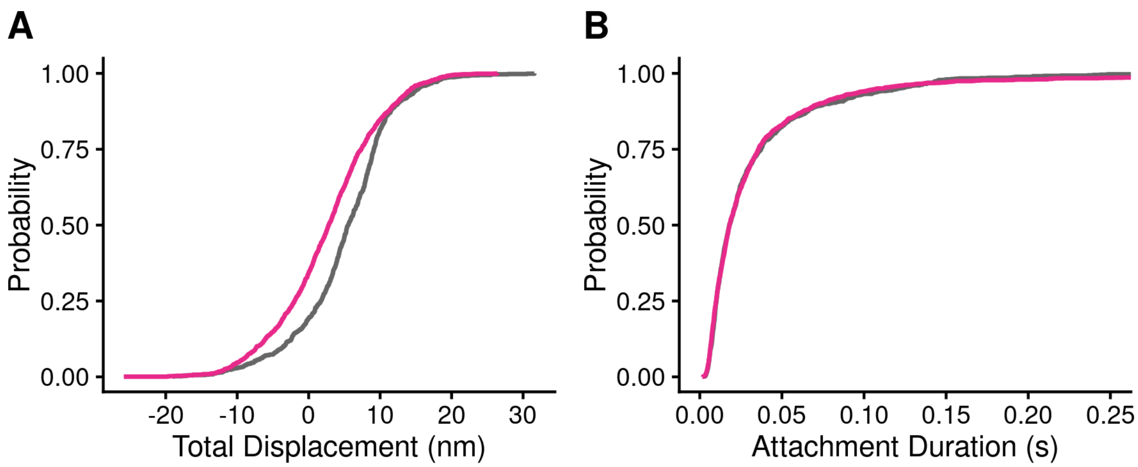


**Supplemental Figure 2: Single molecule optical trapping at 1 mM ATP with and without 10 μM danicamtiv.** Black = DMSO control. Pink = 10 μM danicamtiv. **A)** Cumulative distribution of working stroke displacements. DMSO control has a larger working stroke than myosin treated with 10 μM danicamtiv (5.1 ± 6.8 nm vs. 2.6 ± 7.2 nm, P < 0.001) at 1 mM ATP. **B)** Cumulative distribution of attachment durations. 95% confidence intervals were calculated using bootstrapping methods. There is no different in the detachment rate between treated and DMSO controls (41.4 (-6.6/+6.8) vs. 34.2 (-5.7/+7.1) $s^{-1}$, P = 0.15). Note, the curves overlay. Data also reported in **Table 2** in main text. N = 1755 actomyosin binding events for the control conditions and N = 2107 for 10 μM danicamtiv.

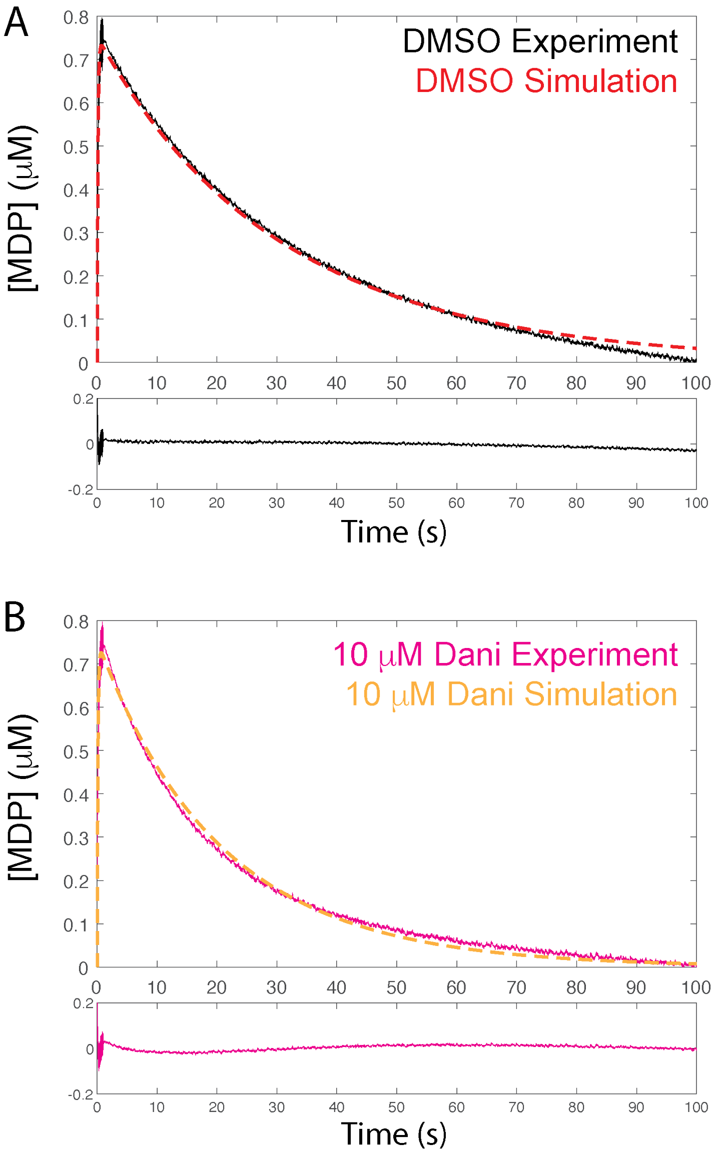


**Supplemental Fig. 3: Fitting single turnover experiments using simulations.** Representative data traces are shown for **A)** DMSO control and **B)** 10 μM danicamtiv. Lower panels show residuals of the fit.

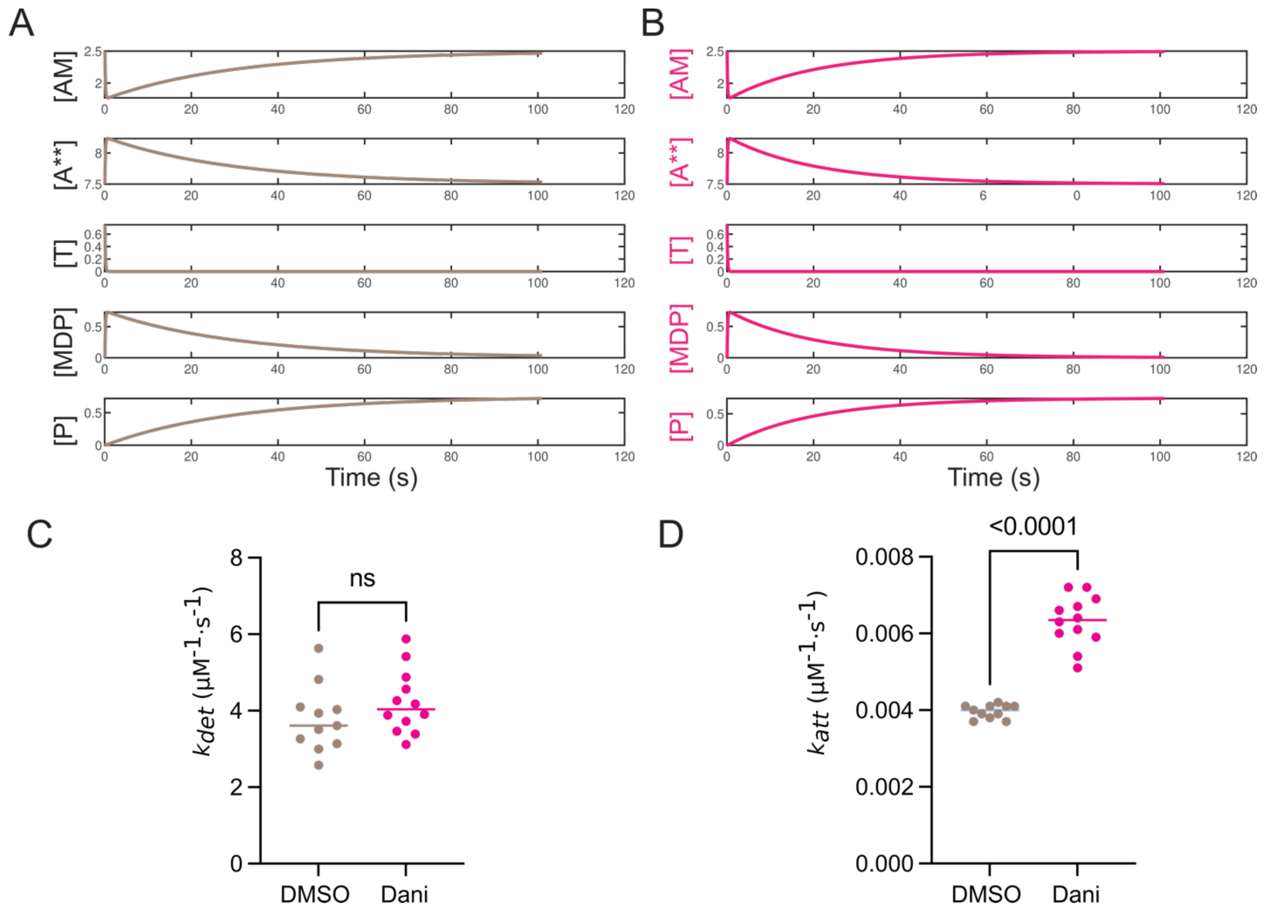


**Supplemental Figure 4: Single turnover simulations.** Calculated chemical species from the best fitted parameters for **A)** DMSO and **B)** 10 µM danicamtiv. AM = actomyosin, A** = unbound actin, T = ATP, MDP = myosin*ADP*Pi, P = phosphate. Best fit parameters for the second-order rates of **C)** actomyosin detachment ($k_{det}$) and **D)** subsequent actomyosin attachment ($k_{att}$). Data is also presented in **Fig. 5**. Each point represents a single fitted transient and bars show the median. Statistical testing was done using a Mann-Whitney test. Note that these are the second-order rate constants that depend on the concentrations of species, not the observed rate constants.

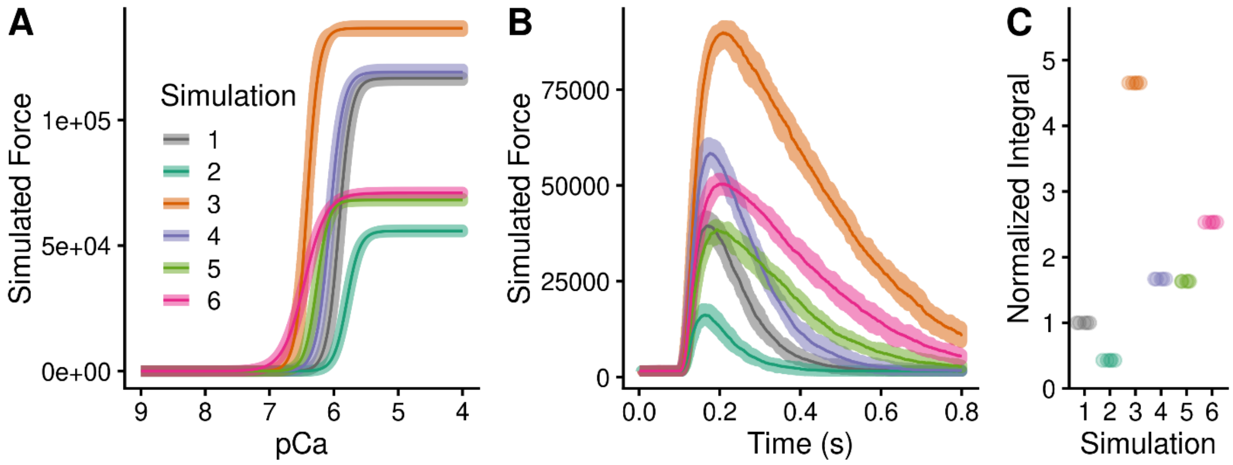


**Supplemental Figure 5: Computational modeling of danicamtiv's effects on muscle contraction. A)** Force calcium-relationship. **B)** Twitch contraction in response to a calcium transient. **C)** Normalized force integrals of the twitch contractions in "B". 5 replicates were conducted, the shaded region shows the range of values, and the solid line shows the mean. The simulations show the effects of:

1) Base model
2) Decreased working stroke (reduced by 1/2 based on optical trapping)
3) Increased actin attachment (increased by 2X based on single turnover measurements)
4) Increased population of active myosin heads (SRX → DRX) (increased by 1.5X based on x-ray diffraction data)
5) Decreased working stroke AND increased actin attachment
6) Decreased working stroke AND increased actin attachment AND increased SRX→DRX
